# Hidden temporal structure of the ongoing task impacts detection strategy and is reflected in pupillary dynamics

**DOI:** 10.1101/2021.11.17.469021

**Authors:** Jennifer Lawlor, Agnès Zagala, Sara Jamali, Yves Boubenec

## Abstract

Estimating temporal regularities in incoming sensory inputs supports optimal decisions in noisy environments. In particular, inferred temporal structure can ease the detection of likely target events. Here we postulated that timely urgency signals can adapt subjects’ decision-making to the ongoing task temporal structure, possibly through neuromodulatory tone. To test this hypothesis, we used an auditory change detection task in which targets followed a block-based temporal contingency, unbeknownst to participants. False alarm occurrences were driven by the distribution of target timings, indicating that participants adapted their behavior to the ongoing temporal structure. Task-evoked pupillary responses were larger for blocks with earliest target timings, and correlated with individual subjects’ behavioral adaptation. Individual pupil responses matched an urgency signal extracted from a decision model fitted to behavior. This work demonstrates that internal temporal expectation can be tracked through pupillary dynamics, suggesting a role of neuromodulatory systems in context-dependent modulation of decision variable dynamics.

## Introduction

Perceptual decisions are often made under uncertainty. Sensory objects are intrinsically noisy, embedded in more complex scenes, and unfold over time. Identifying and reacting to the relevant information in a timely fashion is crucial for interacting successfully with the environment. Temporal expectation is the process of estimating underlying temporal regularities of target events and subsequently making use of those estimates for future behaviorally-advantageous responses. For an ambush predator knowing when to strike is critical (Wearmouth et al. 2014): attacking too early could alert the prey of one’s presence, while waiting too long could lead to being outrun by the prey. In the case of regular events, one can specifically gate detection processes to the expected target timing (Lakatos et al. 2008, 2013). For non-regular events resulting from a stochastic process, decision-making can be biased towards timings when target events become more probable (Tsunoda and Kakei 2008; Oswal, Ogden, and Carpenter 2007; Janssen and Shadlen 2005), indicating that temporal expectation can be factored in perceptual decision-making. Consistent with this idea, temporal dynamics of sensory evidence accumulation during decision-making can change depending on the temporal contingencies of the ongoing task through adaptation of the onset (Teichert, Ferrera, and Grinband 2014) or the duration of the integration process (Ossmy et al. 2013; Piet, Hady, and Brody 2017; Mcwalter and Mcdermott 2018). Understanding how decision-making taps into temporal expectation to optimize behavior is of high importance to complement this picture.

Temporal expectation can be framed as a dynamic signal that biases decision-making at epochs when the relevant signal is likely. Works on speed-accuracy trade-off (SAT), although not studying temporal expectation per se, model decision bias under speed pressure with an urgency component (Murphy, Boonstra, and Nieuwenhuis 2016; Steinemann, O’Connell, and Kelly 2018). In this framework, the urgency component is often directly summed to sensory evidence (Cisek, Puskas, and El-Murr 2009), and then can take the form of either a static offset (van Veen, Krug, and Carter 2008; Hanks, Kiani, and Shadlen 2014) or a linearly growing signal (David Thura and Cisek 2014). Other models incorporate urgency via non-linearly decaying decision boundary (Tajima, Drugowitsch, and Pouget 2016; Palestro et al. 2018). Like the conceptual framework of SAT, temporal expectation has been implemented through an urgency signal which biases the decision variable towards earlier responses (Shinn et al. 2020; Murphy, Boonstra, and Nieuwenhuis 2016; Steinemann, O’Connell, and Kelly 2018; Devine et al. 2019; D. Thura et al. 2012). Intriguingly, recent experimental evidence interprets urgency as a multiplicative factor which amplifies sensory evidence, leading to faster decisions (Yvonne Yau et al. 2021; Y. Yau et al. 2020), as well as impacting evidence-independent motor preparation (Steinemann, O’Connell, and Kelly 2018). Overall, how experimental manipulations of subjects’ temporal expectation impact internal urgency is still a matter of debate.

Neuromodulatory systems play a large role in tuning sensory gain depending on the ongoing context (Jacob and Nienborg 2018; Thiele and Bellgrove 2018; Aston-Jones and Cohen 2005). In line with this idea, modeling works suggest that neuromodulatory signals could mediate a multiplicative urgency-related gain which would adaptively modulate sensory responses (Standage et al. 2011; Niyogi and Wong-Lin 2013). Hence, a compelling proposal would be that neuromodulation can increase sensory gain under speed pressure (Murphy, Boonstra, and Nieuwenhuis 2016), or through the influence of internal temporal expectation. Thus, neuromodulation could enhance detection during a certain time window, at the expense of detection precision. To test this intriguing hypothesis, tracking fluctuations of neuromodulatory tone under different conditions of temporal expectation is a necessary step to determine the role of neuromodulatory systems in adaptive decision-making. Pupil dilation is thought to be under control of the noradrenergic, cholinergic and serotonergic systems (Reimer et al. 2016; Joshi et al. 2016; Murphy et al. 2014; Cazettes et al. 2021) and to reflect a number of cognitive processes (Zhao et al. 2019; Preuschoff, ’t Hart, and Einhäuser 2011; Murphy, Vandekerckhove, and Nieuwenhuis 2014; Krishnamurthy et al. 2017; Knapen and Donner 2014; Keung, Hagen, and Wilson 2019), making pupil size a good candidate for capturing the effects of temporal expectation on neural processes.

Previous works have manipulated speed-accuracy trade-off to characterize how overall urgency affected pupil size (Murphy, Boonstra, and Nieuwenhuis 2016; Steinemann, O’Connell, and Kelly 2018), showing that pupil size increased under speed pressure, possibly due to global gain modulation. Other studies focused on the effect of temporal expectation onto reaction times (Tsunoda and Kakei 2008; Oswal, Ogden, and Carpenter 2007; Shinn et al. 2020) and neural correlates of sensory evidence accumulation (Janssen and Shadlen 2005). However, to date, little is known about the effect of temporal expectation on pupil dynamics. Here we wanted to test how temporal expectation about upcoming target events would be reflected in pupil dynamics throughout trials. For this, we adapted an auditory change detection behavioral task (Boubenec et al. 2017) in which temporal contingencies of target events changed in a block-wise fashion. Trials were divided into 3 blocks defined by their fixed pre-change period (*early, intermediate* and *late* change time blocks). We found that subjects updated their estimation of the change time distribution for each block and that they adapted their behavior accordingly. Pupil size varied in between blocks and correlated with behavioral adaptation of individual subjects. A model of pre-change response with growing urgency captured key behavioral post-change adaptive strategy and reflected block-based pupil dynamics.

## Results

### Auditory change detection task with block-based temporal contingency

We sought to determine if human participants could adapt their perceptual decision strategy to the hidden ongoing temporal structure. For this we modified a previously described change detection task (Boubenec et al. 2017) and manipulated the temporal contingencies of the target events in blocks of trials. Eighteen human participants performed this auditory change detection task. Participants were instructed to report a change in a complex auditory stimulus with a button press without delay. On every trial, a wide-spectrum tone cloud governed by an initial marginal distribution was generated (Figure 1A: example trial). Changes consisted of an increase (of variable size, corresponding to ‘change size’) in tone occurrences in part of the spectrum (‘frequency bin’). Change timings were drawn from an exponential distribution (Figure 1B and Methods: Stimulus) to ensure that participants could not predict target timings on a trial basis. Overall change detection was consistent with previous results reported in (Boubenec et al. 2017) with performance increasing as a function time available to estimate the initial tone cloud statistics until saturation and a dependence on the change size (Figure S1A for detection rate as a function of difficulty and change time and Figure S1B for reaction time as a function of difficulty and change time).

**Figure 1:**
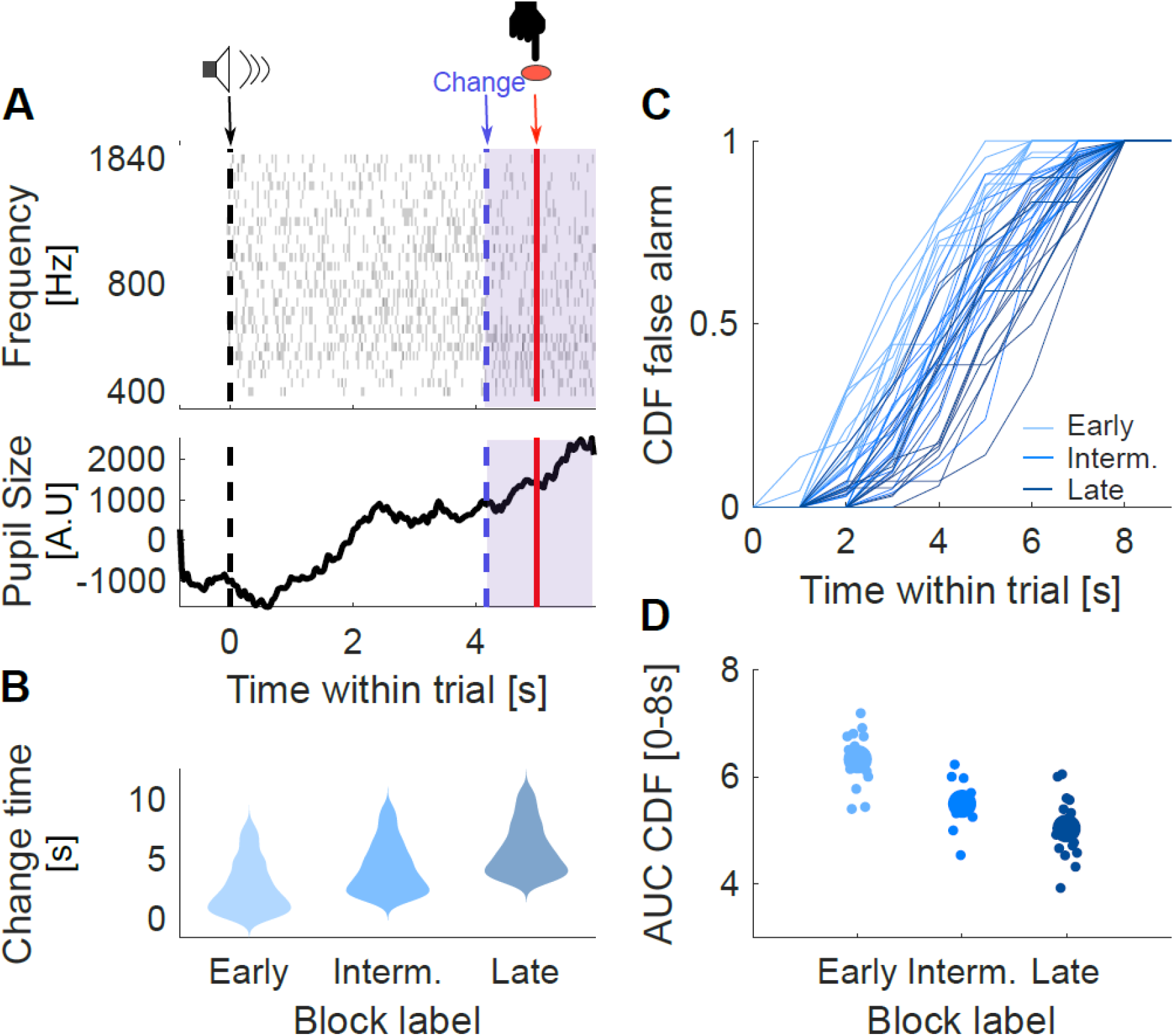
Example trial, experimental block design and false alarm temporal distribution. A. Top panel: Example trial tone cloud as a function of time within the trial. The tone cloud is composed of chords of simultaneous pure tones (grey points, from 400-1840 kHz, duration: 30ms) selected based on a pseudo-random marginal distribution. The change (blue shade) consisted of an increase in tone occurrence probability in one of the frequency bins. In this example the change was presented in the low part of the spectrum. Bottom panel: Pupil size (A.U.) dynamics corresponding to the example trial. Typically, sound onset lead to a small constriction followed by a slow dilation over the course that culminated after the button press. B. Temporal block design: 3 blocks of 120 trials that differed in the start of their change time distribution. *Early* (light blue), *intermediate* (medium blue) and *late* (dark blue) change time distribution, respectively 0 to 8s, 1.5 to 9s, and 3 to 11 s. C. Cumulative distribution (normalized) of false alarm per block as function of time within trial (i.e. from sound onset), individual participant per curve color coded according to temporal block identity. False alarm CDF steepness depends on block design. For the *early* block, false alarms tend to happen earlier than for the *intermediate* and *late* block, respectively. D. Area under the curve for false alarm CDF curve as a function of block. Larger AUC indicates that false alarms distribution is shifted to the left, i.e. false alarms happened earlier (large dot: mean, smaller dot: individual participant). *Early* AUC is significantly higher than *late* AUC (Friedman, p-value = 9.45×10^−5^), suggesting that block temporal identity impacted subjects’ behavior.

The temporal manipulation corresponded to off-setting the change time exponential distribution by a fixed pre-change period for a series of 120 trials (*early* block: 0s, *intermediate* block: 1.5s, *late* block: 3s). Specifically, a change could arise anytime in between 0 and 8s for the *early* block, from 1.5s to 9.5s in the *intermediate* block, and from 3s to 11s for the *late* block (Figure 1B and Methods: Stimulus, *Block temporal structure*). Pupil size was recorded for each participant with the Iscan ETL-200 over the course of a trial (Figure 1A: pupil size example raw trace).

### Temporal expectation modulates early responses

We tested whether the hidden temporal structure of the task could generate differential temporal expectation across blocks. Specifically, we hypothesized that internal expectation about the occurrence of target events at certain times dynamically increases sensory gain, which would lead subjects to respond to stochastic fluctuations in the ongoing stimulus, hence a concomitant increase in false alarm rate. In reaction-time designs (Harun et al. 2020; Johnson et al. 2017; Boubenec et al. 2017), false alarms are defined by a button press prior to the ‘go’ signal. In physiology studies they are referred to as ‘early’ behavioral responses and have been used to link subjects’ strategy and subthreshold stimuli changes (Orsolic et al. 2019). Here, false alarms were of particular interest because, unlike hits and misses, they were not bound by the post-change response window but instead could span the entire range of the pre-change period (up to 11s). Across subjects, false alarm timing distribution shifted as a function of block identity (Figure 1C and D, false alarm timing normalized cumulative distribution [CDF] per subject and normalized CDF Area Under the Curve [AUC], respectively). As predicted, false alarm timings followed the underlying temporal structure of the block; they occurred earlier in the *early* block, than in the *intermediate* and the *late* block (Figure 1C). The mean area under the curve of the normalized cumulative distribution of false alarm timing was significantly larger for the *early*, than the *intermediate* and *late* block (Figure 1D, AUC CDF, Friedman test, p = 9.45×10^−57^).

### Response rates increase as a function of temporal expectation

Our task required subjects to do online detection from the continuous acoustic stream with varying distributions of target timings across blocks. Temporal expectation could then vary differentially in time and across blocks. We therefore tracked temporal dynamics of participants’ response rate. To capture those within the Standard Detection Theory framework we used an instantaneous measure of rates (described in Boubenec et al. 2017 and Methods: Instantaneous rates, criterion and d’ computation). Within each trial, subsegments that did not contain a button press or a change were labeled as correct rejections, leading to an instantaneous false alarm rate. Hit rate was still described as the ratio of hits for trials in which a change was presented to subjects and binned as a function of change time. Both rates were computed with a 1s sliding window and as a function of time within trial (Figure S1C and D, for hit and false alarm rates, respectively). To compare how performance evolved between blocks we restricted our analysis to trials with change timings common to all blocks (from 3 to 8s). Neither block identity, nor time within trial impacted mean instantaneous hit rate over the 3-8s window (Figure S1C, Friedman test, p_block_ = 0.3442, p_time_ = 0.6005). However instantaneous hit rate did vary significantly between 3 and 5.5s as a function of block (Figure S1C, Friedman test, p_block_ = 0.0224, p_time_ = 0.2548). Instantaneous false alarm rate increased with time within trial (Figure S1D Friedman test, ptime = 8.37×10^−7^) but was also clearly modulated by the block identity. Instantaneous false alarm rate was significantly higher for the *early*, than the *intermediate* and *late* block (Figure S1D, Friedman test, p_block_ = 0.0023) over the 3-8s change time window.

### Temporal expectation decreases decision criterion

Increased response rates can have mixed effects on signal detection. Here we postulated that temporal expectation could decrease decision thresholds. Decision criterion captured the subjects’ adaptive strategy. Criterion was lower for the *early* block then the *intermediate* and *late* block (Figure 2A, ANOVA, F(2,26) = 10.98, p_block_=3.4×10^−4^) and also lowered as a function of time within the trial (Figure 2A, ANOVA, F(8,104) = 4.17, p_time_=2.3×10^−4^). d’ decreased as a function of time as instantaneous false alarm rate increased but was not compensated by increase in instantaneous hit rate (Figure 2B, ANOVA, F(8,104) = 3.57, p_time_ = 1.1×10^−3^ and Figure S1C & D for instantaneous hit and fa rate as function of time, respectively). However, there was no significant difference for d’ in between blocks (Figure 2B, ANOVA, F(2,26) = 2.1503, p_block_ = 0.14).

**Figure 2:**
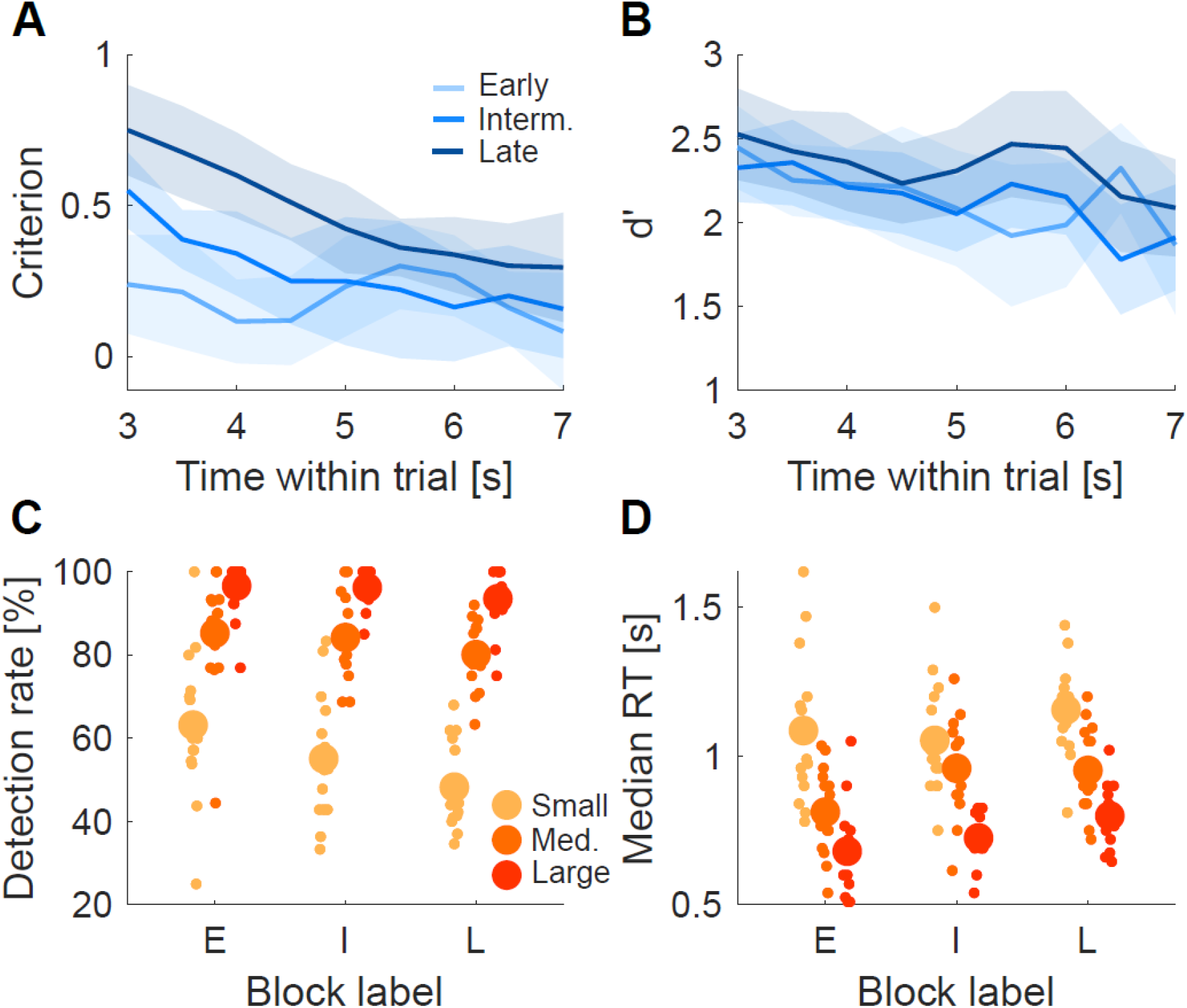
Temporal expectation impacts decision criterion but not sensitivity. A. Mean instantaneous criterion per block varied with time and as a function of block (ANOVA, F(8,104) = 4.16, p_time_ = 0.00023 and ANOVA, F(2,26) = 10.98, p_block_ = 0.00034, respectively). B. Mean instantaneous d’ per block decreased with time but did not vary significantly with blocks (ANOVA, F(8,104) = 3.57, p_time_ = 0.0010 and ANOVA, F(2,26) = 2.15, p_block_ = 0.13, respectively). C. Detection rate (hit rate) per difficulty level (orange color scale), block (x-position) for comparable trials (change time 3 to 8s). Small dots correspond to individual subjects and large dots correspond to the mean performance per condition across subjects. Performance significantly increased with change size for all blocks and decreased with longer pre-change periods (ANOVA, F(2,28) = 117.07, p_diff_ = 3.07×10^−16^ and ANOVA, F(2,28) = 3.93, p_block_ = 0.019, respectively). D. Median reaction time per subject, color convention and dot size are identical to E. Median reaction mirrored performance with larger change sizes, and faster responses for the *early* block (ANOVA, F(2,28) = 59.79, p_diff_ = 7.83×10^−11^ and ANOVA, F(2,28) = 9.33, p_block_ = 7.85×10^−4^, respectively).

### Temporal expectation impacts detection rate and reaction times

Consistent with a lower criterion for the *early* block, the detection rate was highest for that block (Figure 2C, Friedman test average across difficulty, p_block_ = 0.0224), and decreased for the *intermediate* and *late* block. This effect was maximal for smaller change size, with a mean decrease between *early* and *late* close to 15%, compared to a decrease of about 5% for medium and 3% for large changes. Therefore, for equivalent change times, participants were better at detecting smaller changes, suggesting that the amount of evidence necessary to elicit change detection was lower for the *early* block.

Reaction time mirrored detection rate across block and difficulty. Median reaction time, as expected, decreased with change size (Figure 2D, ANOVA, F(2,28) = 59.79, p_size_=7.8e-11). As a function of block identity, median reaction time decreased significantly for medium and large changes (Figure 2D, ANOVA, F(2,28) = 9.33, p_block_ = 7.8×10^−4^).

### Task-evoked pupil response scales with temporal expectation

Here we tested if pupil size reflected participants’ behavioral adaptation to the hidden temporal structure of the task. We restricted our analysis to evoked-pupil responses, e.g. pupil size response after sound onset minus its baseline and normalized per subject (Methods: Pupil diameter traces preprocessing). Task-evoked pupil responses increased as a function of time after sound onset across conditions for most subjects (Figure 3A & C). As predicted, average pupil size for all trials was higher for the *early* block, followed by the *intermediate* and the *late* block (Figure 3A) prior to change detection. This effect is quantified in Figure 3B as the area under the curve for 1.3-2.3s for trials that either contained later changes or button presses (Figure 3B, ANOVA, F(2,28) = 5.54 p_block_ = 9.4×10^−3^). An inverted U-shape relationship has been reported between pupil size, arousal and behavior (McGinley, David, and McCormick 2015; Aston-Jones and Cohen 2005). We expected high arousal states characterized by larger pupil size to have higher false alarm rates. Figure 3C depicts the relationship between trial outcome and pre-change pupil size. Although pupil size for false alarms seemed higher than hits and misses, this effect was not significant (Figure 3C, ANOVA, F(2,28)=0.91, p_outcome_ = 0.41). To ascertain that the reported temporal effect on pupil size is not caused by a larger number of false alarms, we quantified pupil size within block per outcome type in Figure 3D. Even within each outcome, evoked pupil responses increased from *early* to *late* (Figure 3D, ANOVA, F(2,28) = 9.44, p = 0.00073).

**Figure 3:**
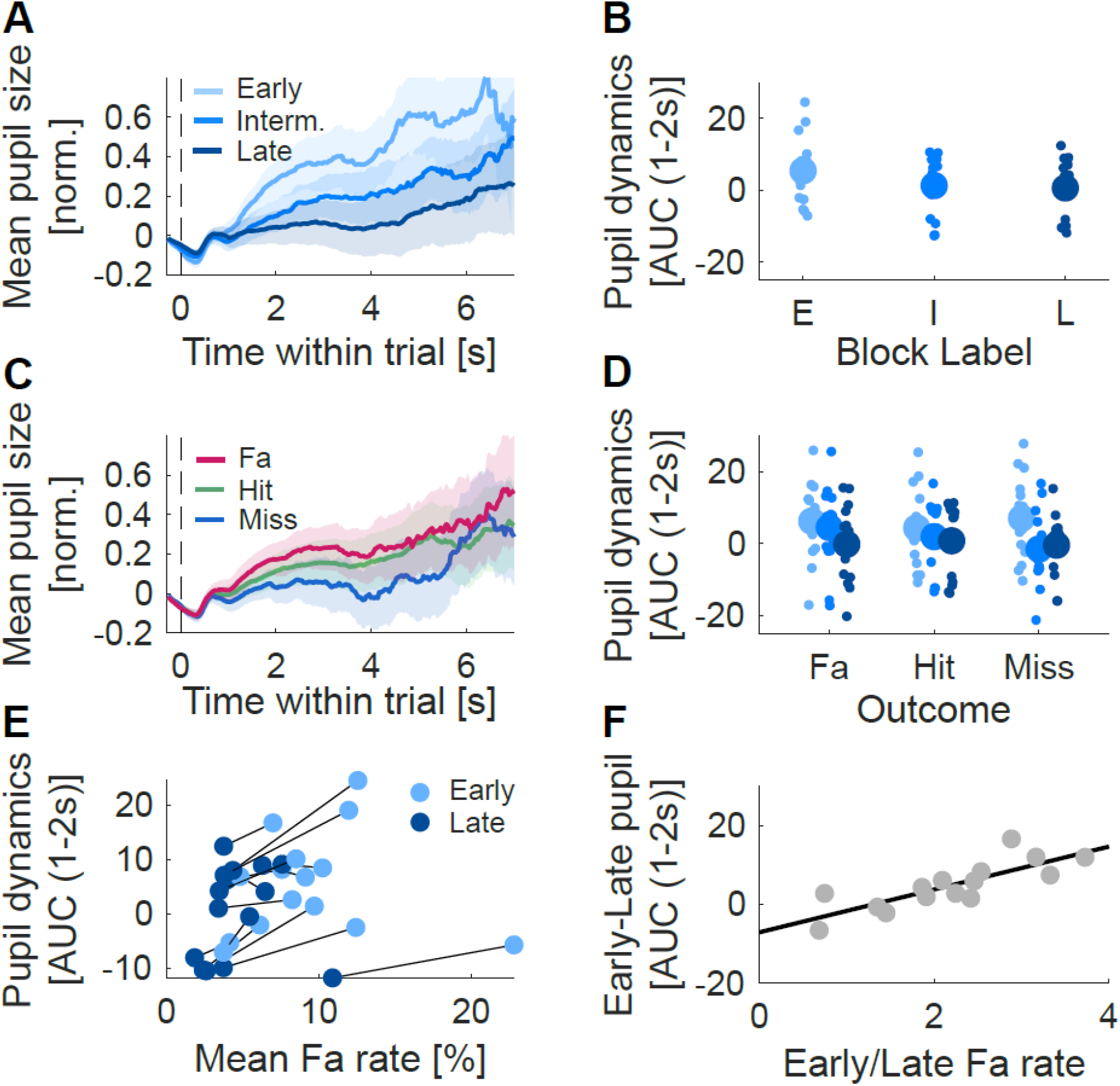
Task-evoked pupil response scales with temporal expectation. A. Mean pre-change task-evoked pupil size per block prior to change (or button press for false alarm) average of all trials over time. Pupil-evoked response was larger for the *early* block, than the *intermediate* and *late* block, as quantified in B. B. Area under the curve for evoked pupil response between 1.3 to 2.3s in trials with change or button press larger than 2.3s per block. Large dot corresponds to mean across subjects (individual: small dot). Task evoked pupil size was largest in the *early* block and scaled with hidden temporal structure (ANOVA, F(2,28) = 5.54 p_block_ = 9.4×10^−3^) C. Mean pre-change task-evoked pupil size per block prior to change per trial outcome. The inverted U-shaped relationship was not significant here (ANOVA, F(2,28)=0.91, p_outcome_ = 0.41). D. Area under the curve for evoked pupil response between 1.3 to 2.3s per block and outcome for each subject (ANOVA, F(2,28) = 9.44, p_block_ =7.34×10^−4^) E. False alarm rate AUC for block *early* and *late* per subject. For most subjects (12 out 15 subjects) false alarm rate decreased with longer change times (*early* higher than *late*) and so does pupil evoked response measured with AUC. F. Linear fit between the ratio of false alarm rate between *early* and *late* block and difference in AUC for the blocks. There was a significant positive relationship between false alarm rate variation and pupil size per subject (linear fit, F-statistic = 25, p-value = 2.43×10^−4^), larger differences in false alarm rate between blocks correlated with larger differences in pupil size.

### Inter-individual differences sensitivity to the task temporal structure are reflected in pupillary dynamics

Previous works reported that pupil size and subject’s overall bias are correlated (de Gee, Knapen, and Donner 2014). The design of our paradigm allowed us to test specifically if pupil size would correlate with subjects’ changes in bias (Figure 2A) that resulted from their adaptation to temporal contingency associated with each block. We regressed the difference in pupil between *early* and *late* blocks against the ratio of false alarms for the same blocks (Figure 3D). We chose the ratio for the false alarm rate to reflect how an individual subject adapted to the temporal manipulation, regardless of their initial bias. Participants’ pupil size and false alarm rate changes across blocks were positively correlated (Figure 3F, Linear Fit, F-statistics = 15.4, p = 0.0018), suggesting that the propension of individuals to make use of temporal regularities scaled the changes in task-evoked pupillary response.

### Ballistic model of urgency captures time-dependent false alarm rates

Psychophysics and modeling works have shown that the effects of speed pressure onto decision-making processes can be explained by an urgency signal. In particular, an additive urgency component can bias the decision variable and ultimately lead to faster reaction times, to the detriment of behavioral accuracy. Here we postulated that temporal expectation could be modeled through an internal urgency signal. Therefore, we predicted that a model of internal urgency could explain the dynamics of false alarms observed across blocks. We first aimed at extracting the characteristics of such an urgency signal and then sought to test that pupil dilations correlated with changes in internal urgency.

For this, we modeled a linearly growing urgency component with a ballistic model (Brown and Heathcote 2008). We focused our modeling effort on reproducing the instantaneous false alarm rates measured in the experimental data. In this model, the urgency component was implemented by a Gaussian distribution of mean slope *ν* and standard deviation *s* (Figure 4A; Methods). Each trial was associated with a drift rate drawn from such a distribution parameterized by *ν* and *s*. Linear drift continued until the response threshold was reached. In this 2-parameter model, within-trial fluctuations were ignored. Randomness in the urgency came from the standard deviation *s* of the drift rate distribution, which resulted in a spread of false alarms over time. We allowed for a single parameter to vary between successive blocks (Methods: Linear Ballistic Urgency model), which implied that we used a total of 4 parameters for fitting the three blocks of different temporal expectation. The model captured the average time-dependent instantaneous false alarm rates across participants (Figure 4B; Supplementary Table 1 for comparison with other models), with a change in slope *ν* between *early* and *late* blocks, and a change in the spread of slopes *s* between *early* and *intermediate* blocks. Linearly growing urgency with a minimal set of parameters can therefore reproduce the temporal profile of false alarm rates under each condition of temporal expectation.

**Figure 4:**
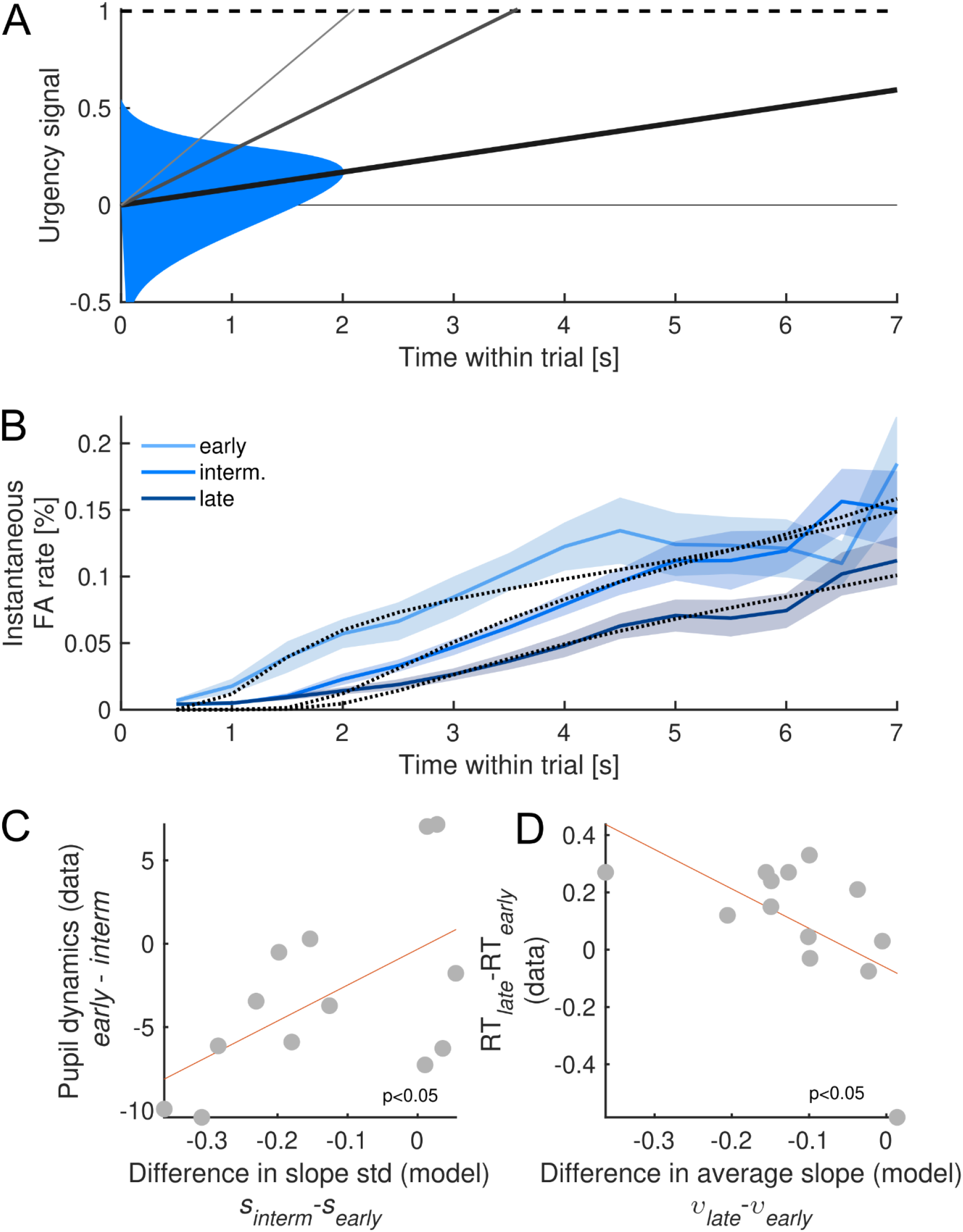
Ballistic model of growing urgency reproduced the effect of temporal expectation. A. Schematic of the ballistic model with linearly growing urgency. Here we depict the Gaussian distribution of urgency slopes for the *intermediate* block. Ballistic trajectories for the mean (black), 1 (dark grey) and 2 (grey) standard deviation slopes are plotted. Response threshold is shown with a dashed line. B. Fit of the ballistic urgency model to the actual instantaneous false alarm rates for each block. The parameters used here were the average over 200 cross-validated fits. C. Change in urgency variance correlated with modulations of pupil size. Larger pupil evoked response in the *early* block correlated with larger variance of the urgency signal in *early* blocks (n = 13; p<0.04). Model parameters were fitted on individual subjects. D. Change in urgency slope correlated with faster reaction times. Gain in reaction times between *early* and *late* blocks negatively correlated with increased slope in *early* blocks (n = 13; p=0.04). Model parameters were fitted on individual subjects.

### Internal urgency correlates with task-evoked pupil responses

Our goal was to correlate internal urgency estimated through the ballistic model with subject-wise modulations of pupil responses by temporal expectation. Hence, we fitted the selected model to individual participants to obtain an individual proxy for internal urgency. We then tested if subject-level changes in task-evoked pupil responses correlated with the urgency signal recovered from the model. We indeed found that subjects with larger modulation of pupillary responses between *early* and *intermediate* blocks showed larger differences in the model between these same blocks (Figure 4C; p=0.04). This finding suggests that pupillary responses correlated with internal urgency.

The urgency signal pushes the decision variable faster towards response threshold, and should lead to faster reaction times. We reasoned that subjects with large differences in urgency slopes between blocks would show a greater reduction of reaction times in *early* blocks. We did find such a negative correlation between the gain in reaction times from the *early* to the *late* block and the difference in the mean urgency slope *ν* fitted in the *early* and *late* blocks (Figure 4D; p=0.04). This is remarkable as the model was fitted only on the false alarm rates, and not on the hit trials used for reaction times. This result indicates that subjects with maximal difference in urgency slopes were showing maximal gain in reaction times.

## Discussion

Detection of behaviorally relevant events in ethological context is a noisy process as sensory signals evolve over time and are embedded in complex scenes. However, event detection can be facilitated by extracting co-varying features in the environment, such as reward contingencies and temporal regularities (Shinn et al. 2020; Ossmy et al. 2013). We aimed at determining if temporal expectation of target events built over multiple trials could inform the detection process and if subsequent changes in subjects’ strategy would be reflected in evoked pupillary dynamics as physiological indicator of neuromodulatory activity. We report that subjects adapted their behavioral strategy to the hidden temporal structure presented in blocks and defined by their change timing distribution. In line with previous reports using visual perceptual decision tasks (Orsolic et al. 2019; Shinn et al. 2020), we found that temporal expectation drove the rate of false alarms throughout trials. Further, we used task-evoked pupil diameter dynamics to determine if neuromodulatory activity could support temporal expectation impact on target detection. Since pupil size increase as a function of subject’s bias (de Gee, Knapen, and Donner 2014) and urgency-scaled pupil signal in SAT paradigms (Murphy, Boonstra, and Nieuwenhuis 2016; Steinemann, O’Connell, and Kelly 2018) have previously been reported, we expected pupil size to be highest in the *early* block.This hypothesis was supported by our data with a higher pupil trace for the *early*, than the *intermediate* and *late* block. More strikingly, subjects’ behavioral adaptation to the hidden temporal structure correlated with changes in subject’s pupillary dynamics across blocks.

### Impact of temporal expectation on decision strategy

An internal urgency signal has been theorized to explain decision-making under speed pressure. Here, we show that a ballistic model with an urgency component fitted to pre-change behavioral responses correlated with subjects’ pupil changes and post-change reaction times. Our study shows that such an internal urgency signal can be generated without instructions from task-related contingencies over multiple trials and can improve detection of events.

The physiological mechanisms that could underlie this time-varying signal are still under investigation. Behavioral effects of speed pressure could originate from 1) evidence-independent motor-preparation (Steinemann, O’Connell, and Kelly 2018), 2) evidence-independent lowering of criterion for action selection (Tajima 2016, Palestro 2018), and 3) neural gain enhancing evidence representation (Yau 2020, Yau 2021, Hawkins Brown 2015). Interestingly, a recent EEG study suggests a hierarchical intertwined implication of an evidence-independent motor component with evidence integration (Steinemann, O’Connell, and Kelly 2018). The dual effect on both the slope *ν* and standard deviation *s* that we reported in our ballistic urgency model could similarly originate from several of those mechanisms. Also, we note that previous models of SAT and temporal expectation did not include trial-to-trial variability (*s* parameter in our ballistic model) in the urgency signal, which was a key ingredient of our modeling approach.

### Neuromodulatory control of urgency reflected by pupillary dynamics

Modeled urgency is proposed to be under neuromodulatory control (Standage et al. 2011; Niyogi and Wong-Lin 2013); (Murphy, Boonstra, and Nieuwenhuis 2016). Pupil dilation reflects the activity of multiple neuromodulatory networks (Reimer et al. 2016; Joshi et al. 2016; Murphy et al. 2014; Cazettes et al. 2021), and those networks in turn support a wide variety of cognitive functions, from attention to learning (Guo, Robert, and Polley 2019; Aston-Jones, Rajkowski, and Cohen 1999; Aston-Jones and Cohen 2005; Janitzky et al. 2015). As such, pupil dynamics has been used as a physiological proxy to investigate the role of neuromodulation in cognitive control. Here, adaptive urgency under temporal contextual pressure correlated with pupil size confirming the implication of neuromodulatory networks in driving time-varying signals related to decision-making. However, how each of those systems impacts decision-making specifically and how their activity interact is still unknown. The source of this physiological effect remains poorly described, and further physiological studies into the neuromodulatory drive of decision-making are needed.

Other subcortical targets activity correlated with time-varying urgency signals such as the globus pallidus in monkeys (Thura and Cisek 2017) and the caudate in humans (Y. Yau et al. 2020). In addition, microstimulation of Superior Colliculus (SC) and Inferior Colliculus (IC) also trigger pupil dilation (Joshi et al. 2016). Recent evidence supports the ‘common drive hypothesis’ where the SC coordinates both saccades and pupil dilation (Wang and Munoz 2021). The SC has also been implicated in the decision criterion generation (Crapse, Lau, and Basso 2018), leading to an intriguing hypothesis: the SC may play a role in mediating a ‘temporal orientation’ response, akin to its traditional role in spatial selection of targets.

### Online tracking of subthreshold events

Natural scenes complexity contrasts with the ease with which humans and animals detect and react to relevant events. The brain’s ability to integrate and monitor sensory inputs over long timescales is key for overcoming this challenge during the decision-making process. For example, natural sounds such as textures can be represented through summary statistics over seconds (McDermott and Simoncelli 2011; McDermott, Schemitsch, and Simoncelli 2013), and identifying relevant behavioral signals in such evolving sensory environments requires building faithful representation of signals over seconds, while still monitoring rapid changes and potential relevant deviations (Mcwalter and Mcdermott 2018; McDermott, Schemitsch, and Simoncelli 2013; Johnson et al. 2017; Boubenec et al. 2017; Devine et al. 2019; Skerritt-Davis and Elhilali 2021). So far, urgency-generated temporal context has not been studied in such situations, and previous works instead used paradigms in which the foreperiod was not informative for the task at hand (Devine et al. 2019; Shinn et al. 2020; Thura et al. 2012). For this reason, we chose a time-varying complex and stochastic stimulus where each time step can be informative of either the pre-change or post-change sound statistics, aiming at mimicking a more ethologically relevant decision-making process in which participants integrated sensory information in both the pre-change and the post-change periods.

Behavioral adaptation to the hidden temporal context led to earlier and increased numbers of false alarms. Interestingly, and because of the stochastic nature of our stimulus, false alarms are likely linked to local changes in the stimulus that resembled changes by chance (Harun et al. 2020). In fact, participants often reported after the task that they perceived changes before false alarms. The shift in subjects’ criterion in the *early* block suggests that participants were more susceptible to report small fluctuations of the pre-change stimulus that resembled the post-change statistics to detect. Unfortunately, our stimulus design did not allow us to specifically identify false alarms that are based on subthreshold fluctuations, and future experiments will be required. Orsolic et al. in 2021 investigated the representation of such subthreshold events in a similar visual reaction time task in mice and used wide-field calcium imaging to describe target selection. They found that the smaller subthreshold events were enhanced in motor regions under increased speed pressure, leading the animals being more susceptible to reporting them. Altogether, this implies that temporal expectation built over multiple trials informs decision-making through urgency related signals that impacts the processing of sensory information.

## Methods

### Experimental set-up

#### Participants

18 human subjects (6 females, 12 males, age range: 22-34 years old) reporting normal hearing and with corrected-to-normal vision participated in this psychophysics experiment. The experimental procedure was approved by the Ethics Committees for Health Sciences at Université Paris Descartes. All participants provided informed consent and were compensated at a rate of 15 euros per hour.

#### Acoustic stimulation

Participants were seated in an acoustically-sealed booth (Industrial Acoustics Company GmbH). Both sound presentation and behavioral responses were controlled using a custom-written software in MATLAB (BAPHY, from the Neural Systems Laboratory, University of Maryland, College Park) usually used for extracellular electrophysiology recordings in animals during behavior but adapted for human psychophysics. Acoustic stimuli were sampled at 100 kHz, converted to an analog signal using an IO board (National instruments, PCIe-6353) and delivered diotically through calibrated high-fidelity headphones (Seinnheiser i380). Acoustic stimuli were presented at 70dB. *Behavioral response* Participants were given a custom-built response box, composed of a single button. Button presses were sent to the IO board (National instruments, PCIe-6353) and are therefore recorded with a sub-millisecond precision.

#### Pupil size recording

Pupil diameter was recorded with the ISCAN reflection tracking system ETL-200 (sampling rate: 1000 Hz) while subjects performed the change detection task. Subjects placed their head on a chin-rest 90cm away from a gray screen (31.6 cd/m^2^) with a small central fixation cross. Luminance was kept constant inside the booth throughout the experiment, and black curtains occluded the window of the acoustic chamber. Before the start of each trial, an 800 ms period of silence was introduced, to estimate pupil diameter baseline before sound onset.

### Stimulus

#### Tone cloud

On every trial, the subjects were presented with a tone cloud ranging from 400 to 1840 Hz, and governed by a pseudo-random marginal distribution. All tone clouds consisted of a train of 30ms chords composed of synchronously presented pure tones. Each chord contained a subset of 26 logarithmically spaced pure tones selected according to the probability of tone occurrence defined by the randomly drawn marginal distribution (Figure 1A: example trial).

#### Change

In this auditory change-detection task, participants were instructed to detect a change consisting of an increase in the tone occurrence probability of part of the spectrum. The stimulus is divided in 4 equally log-spaced frequency bands defined by their initial marginal distribution. At the change time, the initial marginal distribution is modified with an increase of probability of variable size (corresponding to the change size) for one of the frequency bands. The increase in local power is compensated by an equivalent decrease in the other bands to maintain global loudness. Change sizes were limited to 3: 60%, 95% and 130%, corresponding to a small, medium and large change. Change time was drawn from an exponential distribution, bounded by blocks’ temporal structure from 0 to 11s.

Participants had 2 seconds to detect and react to the change. A button press before the change occurrence was labelled a false alarm, after the change and within the 2s response window, a hit and if there was no button press 2s after the change, a miss.

#### Block temporal structure

Unbeknownst to the subjects, change time blocks were introduced. In each of these blocks, a change could arise only after a fixed pre-change period (*early*: 0s, *intermediate*: 1.5s and *late*: 3s from trial onset, Figure 1B). Blocks consisted of 120 successive trials and no indication of a block beginning or end was given to the participants.

### Procedure

#### Instructions

Each participant received the same set of instructions, mainly that they were going to perform a change detection task, and that each trial contained a change.

#### Training

Instructions were purposefully vague about the nature and or timing of the change. We ensured that participants could perform the task reliably with a training session of approximately 40 trials of the largest change size (130%). Only participants that reached a 40% hit rate were allowed to start the main experiment.

#### Trial structure

Participants were asked to maintain fixation of a cross placed in the center of the screen during trials. Feedback was given at the end of every trial, in the form of a check (‘v’) for correct trials (hits) and a cross (‘x’) for incorrect trials (false alarms and misses). The fixation cross and the feedback objects were the same size and color (white over a grey background) ensuring a constant luminance over the entire experiment. Feedback was displayed after the button press, and subjects were instructed to preferentially blink during feedback presentation. After being displayed for 1s feedback was replaced by the fixation cross for another second before the start of the next trial. Each subject performed a total of 360 trials, divided into 3 blocks of 120 trials each. Block presentation order was randomized across subjects. The experiment lasted about 1 hour and 30 minutes with two 5 min breaks that did not coincide with the beginning or end of a block.

### Pupil diameter traces preprocessing

3 subjects were excluded from further analysis as their pupil diameter was not recorded properly (software thresholds were not respected, leading to only one value for pupil diameter), leaving a total of 15 subjects for the analysis.

#### Blink removal

Blinks defined as points outside 1.5 standard deviation of the mean pupil trace (over the entire session) were replaced by NaNs over an asymmetric window spanning 350ms (150ms prior, 200ms after identified blink event).

#### Filtering

The pupil time series were first low-pass filtered (third order Butterworth, cutoff, 10 Hz, (de Gee, Knapen, and Donner 2014).

#### Task-evoked pupil response

To investigate the effect of temporal expectation on pupil size dynamics within trials, we focused on task-evoked pupil response by subtracting the mean pupil diameter estimated during the baseline period from the corresponding trial. In addition, to allow for across subjects comparison (Iscan output is in arbitrary units), pupil traces were divided by each subject’s standard deviation leading to comparable pupil size distribution.

### Behavioral analysis

We quantified participants’ ability to detect changes in different conditions (block-based temporal and change size) using time-dependent hit, false alarm rates, and d’ and criterion estimation developed in (Boubenec et al. 2017). This allowed us to compare subjects’ responses at matching points in time bins without having to restrict our analysis to trials with matching change time. Block-based comparison of performance or reaction time is done for change timings that are common across all blocks [3-8s], and for which the estimation of baseline statistics has saturated.

### Instantaneous rates, criterion and d’ computation

#### Instantaneous hit rate

HR(t) was computed as the fraction of correct change detections, in relation to the number of trials with changes occurring over a 1s window centered around time t. For this specific paradigm, HR(t) was computed for 14 overlapping time bins from 0s to 7.5s in 0.5s increments. For each segment, the hit rate is: HR(t) = H(t) / (H(t)+M(t)), H(t) corresponds to the number of hits and M(t) to the number of misses for change timings occurring within this time bin.

#### Instantaneous false alarm rate

Similarly to the d’ time-dependent measure in (Boubenec et al. 2017), trial segments before the change were used as correct rejections. FAR(t) was computed for 14 overlapping time bins from 0s to 7.5s in 0.5s increments. For each segment, the false alarm rate is: FAR(t) = FA(t) / (FA(t)+CR(t)), FA(t) corresponds to the number of false alarms occurring within this time bin (timing of the button press for a false alarm) and CR(t) is the number of trials where the change appears after the end of the time bin.

#### Instantaneous d’

We used the classical approximation d’(t) = Z(HR(t)) - Z(FAR(t)) to compute participants’ sensitivity as a function of time per condition. Z(p) is the inverse of the Gaussian cumulative distribution function (CDF). HR(t) is the hit rate as a function of time since stimulus onset, described above. FAR(t) is the false alarm rate as a function of time since stimulus onset, described above.

#### Instantaneous criterion

We used the classical approximation b(t) = (Z(HR(t))+Z(FAR(t)) / 2 to compute participants’ bias as a function of time per condition.

### Reaction times

Reaction times were obtained by subtracting the change time from the response time hit trials. For each condition, the distribution of reaction times was assembled and the median reaction time computed per subject. Figures usually present the mean reaction time across subjects with individual medians displayed with a smaller dot (Figure 2F).

### Pupil dynamics analysis

Pupil dynamics analysis was restricted to phasic response locked to stimulus onset and corrected by the silent pre-onset stimulus baseline (Steinemann, O’Connell, and Kelly 2018; de Gee et al. 2020; Zhao et al. 2019). To investigate the effect of temporal expectation on task-evoked pupil response we quantified the pupil dynamics as the area under the curve for comparable pre-change time points between 1.3 and 2.3s. We picked this specific window to 1) maximize the number of trials per conditions, since trials containing change before 2s are removed for the analysis, and 2) let the pupil diameter recover from the initial onset-related constriction (Figure 3A and B).

### Statistical analysis

We tested each measure for normality using the Shapiro-Wilk test. If data was reported as normal (such as pupil dynamics AUC) we conducted within subject repeated measures ANOVAs, to determine significance of the temporal manipulation of our measures. For data that was not normally distributed (such as performance) we conducted significance testing using Friedman tests. Only reaction times distribution was modified to allow for parametric testing by using the log. All data analysis and statistical analysis was performed using custom-built script in MATLAB.

### Linear Ballistic Urgency model

We postulated that changes in responsiveness across blocks came from changes in subject internal urgency. For extracting a measure of internal urgency, we adapted a classical model of decision-making called Linear Ballistic Accumulator (LBA, Brown and Heathcote 2008). This model is classically used for fitting response time distributions under different conditions of speed-accuracy trade-off (Heitz and Schall 2012; Ho et al. 2012). LBA models assume a linear accumulation of evidence towards a decision, with an absence of randomness within trial. Variance in reaction times arises from the variance in the drift rate distribution. In the model used here, drift rates are drawn from a Gausian distribution of mean drift rate *ν*, and standard deviation *s*. Here we substituted the accumulator variable of the LBA with an urgency variable, with identical parameterization *ν* the mean urgency drift rate and *s* the standard deviation of the drift rate distribution (Figure 4A; (Adam, Bays, and Husain 2012). We chose to fit the instantaneous false alarm rates which is a metric independent of trial number, but instead reflects the propensity to respond at every point in time. Despite their conciseness, LBA models bring similar conclusions than accumulator models which include within-trial randomness (Usher and McClelland 2001; Donkin et al. 2011). This makes LBA-type models an appealing modeling choice for working with a reduced set of parameters.

Close inspection of false alarm rates suggests that there are quantitative differences between the *early* block, and the other two blocks (Figure 2B and 4B). False alarms in the *early* block showed an early take-off (before 2s), which was absent in the *intermediate* and *late* blocks for which false alarms increased no earlier than 2s. On the contrary, the difference between *intermediate* and *late* blocks was in their average slope. We expected this quantitative difference to impact the ballistic model. We therefore plotted false alarm timing with reciprobit plots (Noorani and Carpenter 2016), so that distinct effects on the *ν* or *s* parameters would be isolated (Figure S2). Reciprobit plots for the *intermediate* and *late* false alarm timings showed an horizontal shift of the response times between *intermediate* and *late* blocks, which indicates a change in the mean drift rate *ν*.

Based on these observations, we hypothesized that one single parameter (*ν* or *s*) varied between blocks of adjacent expectation (*early* to *intermediate, intermediate* to *late*). This led to 4 different alternative models. We fitted these 4 different models (supplementary table S1), and found that model (n°4) described the best our data. Model n°4 postulated a change of the standard deviation of the drift rate distribution for the *early* block, and a change of mean urgency drift rate between the *intermediate* and *late* blocks, which matches the observations of the reciprobit plots.

**Supplementary Table 1:** Summary table of the goodness-of-fit for the different tested models. Each model has 4 free parameters (a-d), with each condition described by two parameters *ν* and *s*, with one parameter changing between adjacent conditions.

| Block | Early |  | Intermediate |  | Late |  | MSE (10<sup>-3</sup>) |
| --- | --- | --- | --- | --- | --- | --- | --- |
| Parameter<br>s | v | s | v | s | v | s |  |
| Model 1 | a | b | a | c | a | d | 0.6943 +-0.39 |
| Model 2 | a | b | c | b | d | b | 0.808 +-0.42 |
| Model 3 | a | b | c | b | c | d | 0.77 +-0.42 |
| Model 4 | a | b | a | c | d | c | **0.6469** +-0.40 |

### Procedure for fitting and evaluating models

Procedure for testing the goodness-of-fit of the four alternative models was as follows: we performed 200 cross-validations in which we fitted the four free parameters on the average instantaneous false alarm rates obtained over 8 randomly drawn subjects. We then computed the MSE between the model predictions and the average over the 7 left-out subjects. This splitting procedure assumed that both groups would share the same inter-individual variability. We used the *fminsearch* algorithm from MATLAB which is based on the Nelder-Mead simplex methods. To prevent incorrect fit due to local minima, every cross-validation was averaged over 20 fits done with different randomly chosen starting seeds drawn between 0 and 1.

For a drawn urgency drift *ν*_0_, false alarm timing will be 1/*ν*_0_. Following this logic, distribution of false alarm timing for a set of parameters *ν* and *s* can be computed as:

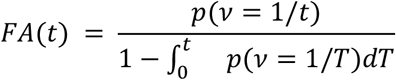

For single subject fit, we initialized the fitting procedure with the average parameters recovered across the 200 cross-validations performed on the average data.

**Supplementary Figure S1.**
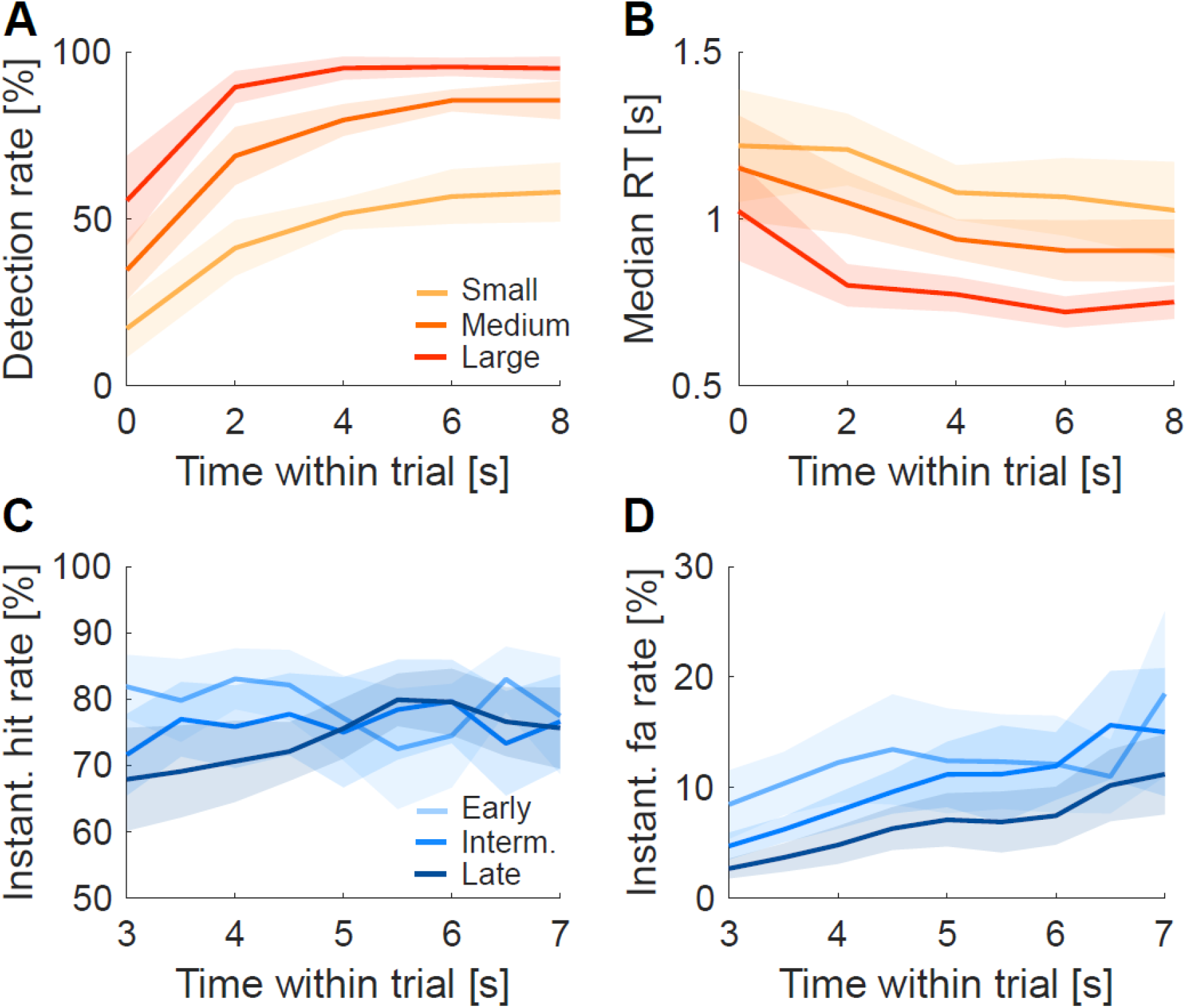
A. Detection rate (as hit rate) increased as a function of time within the trial and difficulty level (as change size), reproducing results described in Boubenec et al. 2017. B. Median reaction time decreased as function of time within the trial and difficulty level (as change size), mirrorring detection rate in A. C. Mean instantaneous hit rate (shaded area: SEM) as a function of time within trial for comparable change timings across blocks (3 to 8 s), color code corresponds to block identity. Mean instantaneous hit rate was not impacted over the 3-8s change time window by time within trial or block identity (Friedman test, p_block_ = 0.3442, p_time_ = 0.6005). However instantaneous hit rate did vary significantly between 3 and 5.5s as a function of block (Friedman test, p_block_ = 0.0224, p_time_ = 0.2548). D. Mean instantaneous false alarm rate (shaded area: SEM) as a function of time within trial for comparable trials across blocks (3 to 8 s), color code corresponds to block identity. False alarm rate was highest for the *early* block and decreased significantly for the *intermediate* and *late* block (ANOVA, F(2,26) = 8.07, p_block_ = 0.0017) and increases significantly as a function of time (ANOVA, F(8,104) = 9.62, p_time_ = 4.5×10^−10^).

**Supplementary Figure S2:**
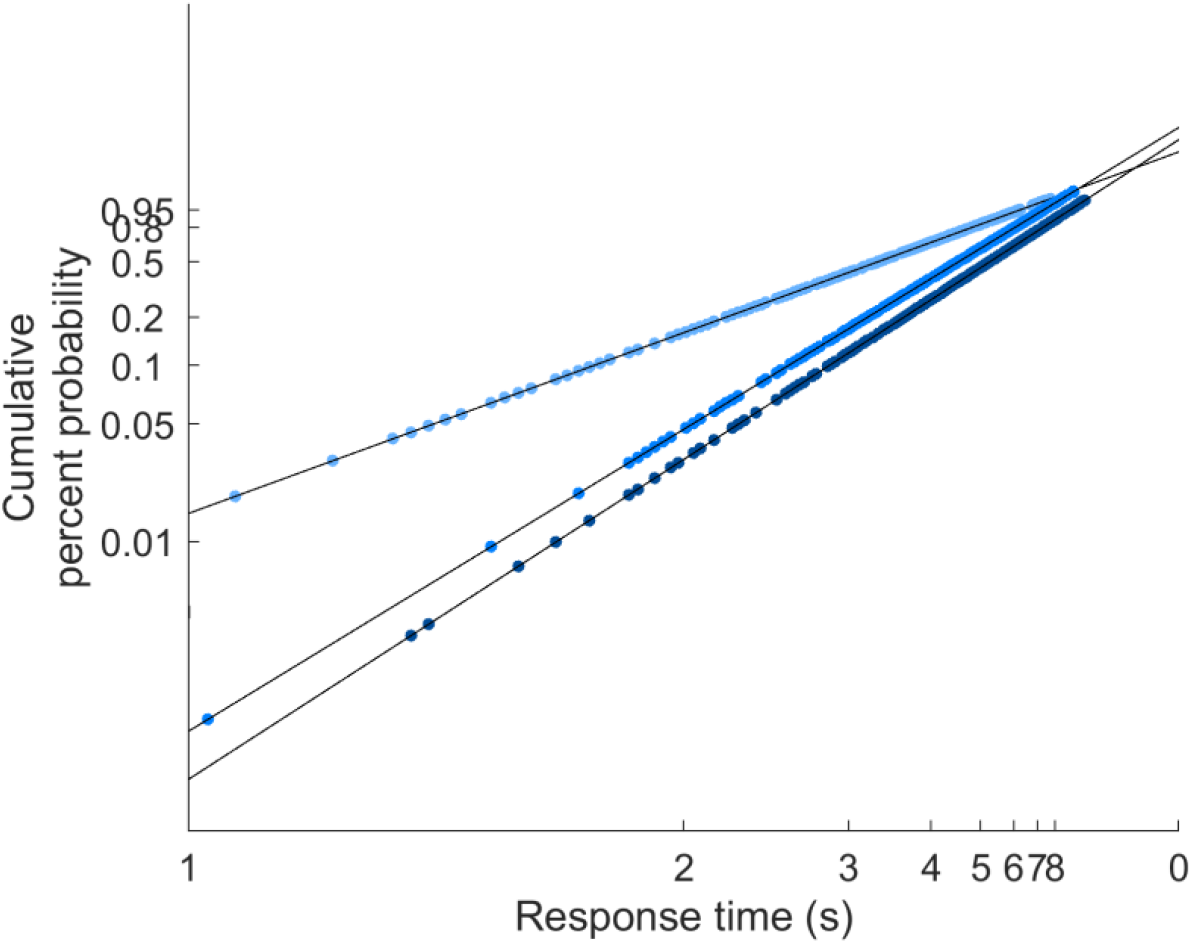
Reciprobit plots of the false alarm response times.

## Author Contribution

J.L. and Y.B. designed the experiment, analysed, and wrote the manuscript.

A.Z. collected the data and started the analysis process.

S.J. helped with data analysis.

## Acknowledgements

We thank Dr Shihab Shamma, Dr Daeyeol Lee, Dr Ninad Kothari, Dr Kate Allen and Clarice Diebold for their thoughtful comments and suggestions on our manuscript. Our thanks to Dr Bernhard Englitz for his crucial work on the psychophysical task and fruitful guidance. We especially thank Dr Célian Bimbard for endless discussion of this topic during and after data collection.

